# Robustness of the ferret model for influenza risk assessment studies: a cross-laboratory exercise

**DOI:** 10.1101/2022.04.02.486825

**Authors:** Jessica A Belser, Eric HY Lau, Wendy Barclay, Ian G. Barr, Hualan Chen, Ron AM Fouchier, Masato Hatta, Sander Herfst, Yoshihiro Kawaoka, Seema S Lakdawala, Leo Yi Yang Lee, Gabriele Neumann, Malik Peiris, Daniel R Perez, Charles Russell, Kanta Subbarao, Troy C Sutton, Richard J Webby, Huanliang Yang, Hui-Ling Yen, Working group on the standardization of the ferret model for influenza risk assessment

## Abstract

Ferrets represent the preferred animal model for assessing the transmission potential of newly emerged zoonotic influenza viruses. However, heterogeneity among established experimental protocols and facilities across different laboratories may lead to variable results, complicating interpretation of transmission experimental data. Between 2018-2020, a global exercise was conducted by 11 participating laboratories to assess the range of variation in ferret transmission experiments using two common stock H1N1 influenza viruses that possess different transmission characteristics in ferrets. Inoculation route, dose, and volume were standardized, and all participating laboratories followed the same experimental conditions for respiratory droplet transmission, including a strict 1:1 donor:contact ratio. Additional host and environmental parameters likely to affect influenza transmission kinetics were monitored throughout. Overall transmission outcomes for both viruses across 11 laboratories were concordant, suggesting the robustness of the ferret model for zoonotic influenza risk assessment. To attain high confidence in identifying zoonotic influenza viruses with moderate-to-high or low transmissibility, our analyses support that as few as three but as many as five laboratories, respectively, would need to independently perform viral transmission experiments with concordant results. This exercise facilitates the development of a more homogenous protocol for ferret transmission experiments that are employed for the purposes of risk assessment.

## Introduction

Pandemic influenza viruses may arise through interspecies transmission events of animal influenza viruses. Assessing the human-to-human transmission potential of animal influenza viruses that cause spillover infections in humans is essential for pandemic risk assessment. Ferrets have been used as a surrogate model for investigating the transmission potential of influenza viruses in humans, as they are naturally susceptible to infection with human and zoonotic influenza viruses, exhibit clinical signs during infection which closely resemble those of humans, and support influenza virus transmission via similar modes as humans. In particular, the respiratory droplet transmissibility of a specific influenza strain among ferrets often correlates with its transmission potential in humans (1). As such, ferrets are commonly used for assessing the pandemic potential of newly emerged zoonotic influenza viruses, and data from these experiments inform formal risk assessment rubrics established by the WHO and CDC (2, 3).

The transmission potential of influenza viruses is determined by multiple viral, host, and environmental parameters. As the ferret model becomes commonly employed in laboratories worldwide, there is an underappreciated heterogeneity among established experimental protocols and facility setups across different laboratories, which may lead to variable results between transmission experiments performed (4). Some of these variables, such as the dose, volume, and route of inoculation and animal age, have been confirmed to affect the kinetics of virus infection, replication, and transmission in the ferret model (5–7). However, the impact of other parameters, such as virus propagation procedures, caging designs, airflow directionality and number of air exchanges, and environmental conditions such as relative humidity, is largely unknown. Consequently, interpretation of results from ferret transmission experiments can represent a challenge when comparing data generated from multiple laboratories, even when the same virus strain or subtype is being investigated (8). Considering the statistical limitations on small sample sizes in ferret experiments, and high potential for strain-specific variability, investigators often assess the pandemic potential of emerging virus subtypes as an aggregate of multiple viruses tested (9–11). As many public health efforts require cross-laboratory risk assessment studies for newly emerged zoonotic influenza viruses (12) and antiviral efficacy studies aiming to block influenza transmission between ferrets (13), a greater understanding of variability in transmission results obtained between independent groups is critical.

To assess the variability of ferret transmission results across laboratories under established protocols, we performed a global exercise using two common stock influenza viruses that possess different transmission characteristics in ferrets. Eleven independent laboratories inoculated ferrets with these stock viruses under uniform conditions; parameters known to affect influenza transmission kinetics were controlled in the experimental protocols while other potential parameters were carefully monitored and recorded, both prior to and during the transmission experiments. All aggregated data from these experiments were blinded and analyzed by an independent statistician. To inform future risk assessment activities, the confidence of drawing conclusions on virus transmissibility with concordant or discordant outcomes from multiple laboratories was also investigated. By assessing the range of variation present among ferret transmission experiments performed under established experimental protocols, this global exercise provides helpful guidance for data interpretation when cross-laboratory results are to be compared. The relatively concordant transmission results across 11 laboratories suggest that the ferret model is highly robust for influenza pandemic risk assessment studies under the semi-standardized conditions employed here. Furthermore, analyses investigating the role of host and environmental parameters as they contribute to virus transmission kinetics and outcomes is valuable for both current risk assessment activities and for evaluation of countermeasures to block influenza transmission.

## Results

### Transmissibility of human A(H1N1)pdm09 virus

To evaluate potential heterogeneity in the transmission results between 11 laboratories, we first compared the transmissibility of a cell-grown isolate of the A(H1N1)pdm09 virus A/California/7/2009 (Cal/09), representative of early 2009 pandemic isolates and anticipated to exhibit moderate to high respiratory droplet transmissibility (14–17). Transmissibility was evaluated with 4 donor:contact pairs at a 1:1 ratio in each laboratory. Transmission to exposed respiratory droplet contact ferrets was defined by detection of infectious virus or seroconversion to the homologous virus in post-exposure sera. Following establishment of contact with donor ferrets 24 hours post-inoculation, detection of infectious virus and seroconversion in contacts was observed in 10/11 and 11/11 laboratories, respectively, with the reported transmission frequency ranging from 50-100% (Table 1). One out of 11 laboratories determined viral loads in nasal swabs and throat swabs (Group F, with throat swab viral loads used for subsequent analysis), while the other laboratories determined viral loads in nasal washes. Employing both virological and serological results, by Fisher’s exact test of homogeneity, there was no significant difference in the transmission outcomes across labs with this virus (p=0.797). Collectively, infectious virus was detected from the nasal wash or throat swabs of 72.7% (32/44) exposed contacts, and seroconversion of contact ferrets to homologous virus was detected from 79.5% (35/44) of exposed contacts.

**Table 1.** Summary of virus transmissibility results from all laboratories.

| Group | A(H1N1)pdm09 virus,<br>A/California/7/2009 |  |  | A(H1N1) avian influenza virus,<br>A/ruddy turnstone/Delaware/300/20/2009 |  |  |
| --- | --- | --- | --- | --- | --- | --- |
|  | Viral load of<br>inoculated<br>donors (area<br>under the<br>curve)<sup>a</sup> | Transmission to aerosol<br>contacts |  | Viral load of<br>inoculated<br>donors (area<br>under the<br>curve)<sup>a</sup> | Transmission to aerosol<br>contacts |  |
|  |  | Virus<br>detection<sup>b</sup> | Sero-<br>conversion<sup>c</sup> |  | Virus<br>detection<sup>b</sup> | Sero-<br>conversion<sup>c</sup> |
| A | 6.51±0.49 | 3/4 | 3/4 | 4.28±0.35 | 0/4 | 0/4 |
| B | 5.70±0.42 | 4/4 | 4/4 | 4.25±0.39 | 1/4 | 1/4 |
| C | 5.30±0.78 | 4/4 | 4/4 | 5.10±0.11 | 1/4 | 1/4 |
| D | 6.86±0.40 | 2/4 | 2/4 | 5.73±0.21 | 0/4 | 0/4 |
| E | 5.53±0.32 | 3/4 | 3/4 | 4.43±0.56 | 0/4 | 0/4 |
| F | 5.77±0.60 | 3/4 | 3/4 | 5.34±0.60 | 0/4 | 0/3 |
| G | 6.57±0.06 | 2/4 | 2/4 | 6.48±0.37 | 0/4 | 0/4 |
| H | 5.82±0.43 | 0/4 | 3/4 | 4.72±0.31 | 0/4 | 0/4 |
| I | 6.48±0.80 | 3/4 | 3/4 | 6.24±0.31 | 0/4 | 0/4 |
| J | 5.62±0.54 | 4/4 | 4/4 | 4.92±0.39 | 1/4 | 1/4 |
| K | 4.07±0.72 | 4/4 | 4/4 | 4.15±0.57 | 3/4 | 3/4 |

<sup>a</sup>Area under the curve (AUC) of viral titers from inoculated ferrets (following normalization of infectious units to TCID<sub>50</sub>), expressed as log<sub>10</sub> mean ± standard deviation. <sup>b</sup>Number of contact ferrets with infectious virus detected in respiratory specimens between 1-13 days post-contact (p.c.). <sup>c</sup>Number of contact ferrets that seroconverted to the exposed virus at the end of the study using hemagglutination inhibition assay.

To allow comparison of the effect of viral load on transmissibility, viral titer units from nasal wash/throat swab samples (inclusive of TCID_50_, PFU, and EID_50_ units, Supplemental Figures 1-2) were normalized to TCID_50_ units (Figure 1), employing strain-specific conversions prior to analyses (Supplemental Table 1). From the inoculated donor ferrets, the peak viral titers detected in the nasal washes or throat swabs were at 5.72 ± 0.95, mean ± SD log_10_ TCID_50_/mL after normalization, with the peak titers detected from 95.5% (42/44) of donors on the first sampling time point on 1 or 2 days post-inoculation (dpi), followed by a decline of infectious titer over time (Figure 1A). Area under the curve (AUC) after normalization was calculated to approximate total viral load shed by the Cal/09-inoculated donors, with a mean ± SD log_10_ AUC of 5.84 ± 0.89.

**Figure 1.**
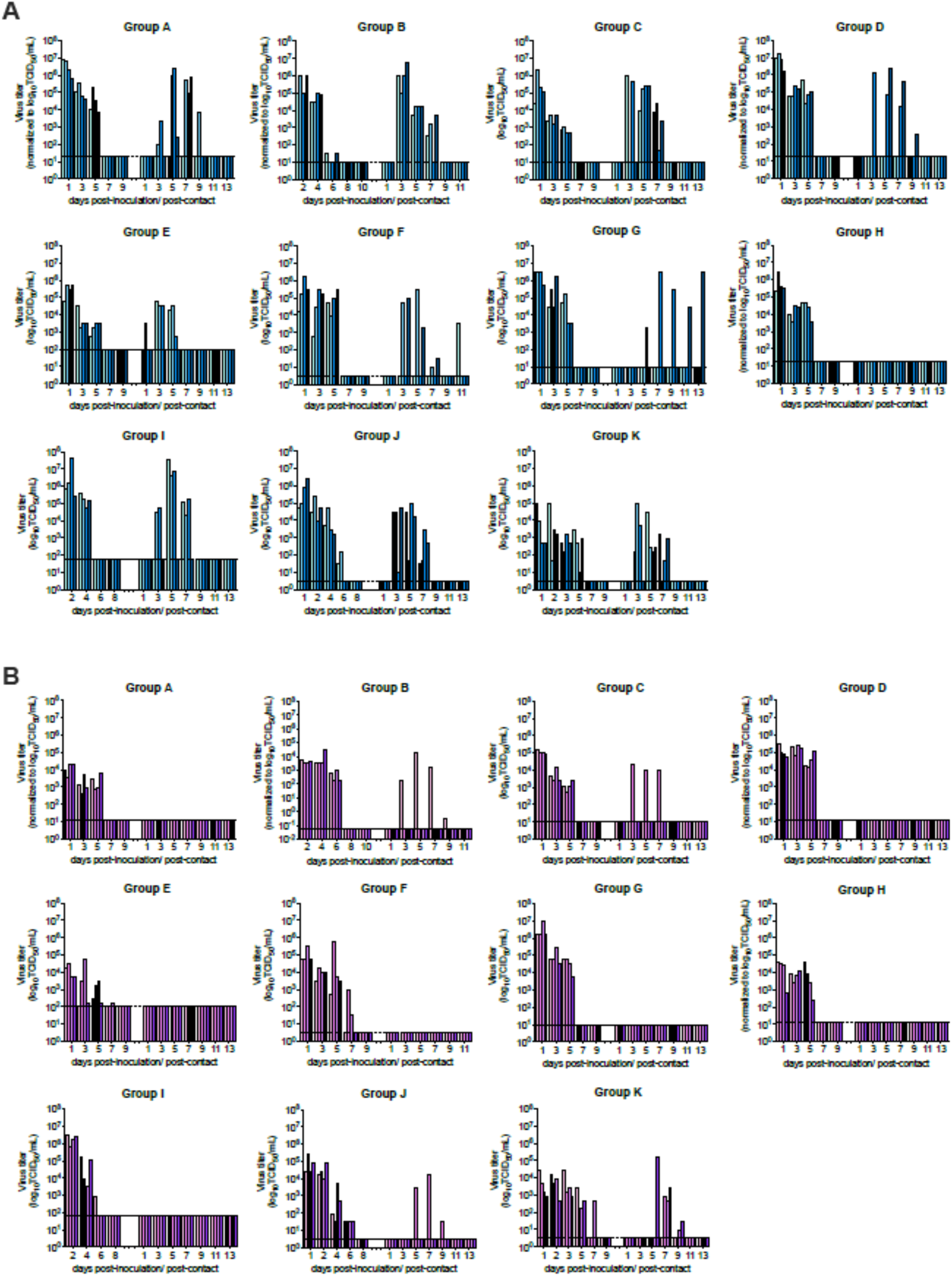
Transmission kinetics of A(H1N1) viruses in ferrets. **A**, normalized viral loads of donors (left bars) and aerosol contact ferrets (right bars) after inoculation or exposure to A(H1N1)pdm09 virus Cal/09. **B**, normalized viral loads of donors (left bars) and aerosol contact ferrets (right bars) after inoculation or exposure to avian H1N1 virus ruddy turnstone/09. Nasal washes (all groups except Group F) or throat swabs (Group F) were sampled to determine infectious viral loads which were normalized to log_10_ TCID_50_/mL. Each bar represents individual ferrets. Limit of detection is indicated with a dashed line.

Next, to evaluate the transmission efficiency, the serial interval (first detection of viral shedding in contacts post-exposure from specimens collected every-other-day) was calculated for each infected contact ferret. The serial interval was 1 day for 3.1% (1/32) of the Cal/09 infected contact ferrets, followed by 3 days for 68.8% (22/32), 5 days for 21.9% (7/32), and 11 days for 6.3% (2/32), with a median serial interval of 3 days post-contact. Peak viral titers detected in the contact nasal washes or throat swabs were at 5.41 ± 1.06 mean ± SD log_10_ TCID_50_/mL after normalization, with peak titers detected from 50% (16/32) and 34.4% (11/32) infected contacts on 3 dpi and 5 dpi, respectively. Altogether, the AUC for Cal/09 infected contact ferrets was 5.75 ± 1.05, comparable to the Cal/09 virus-inoculated donors (Mann-Whitney test, p=0.6547) (Figure 2).

**Figure 2.**
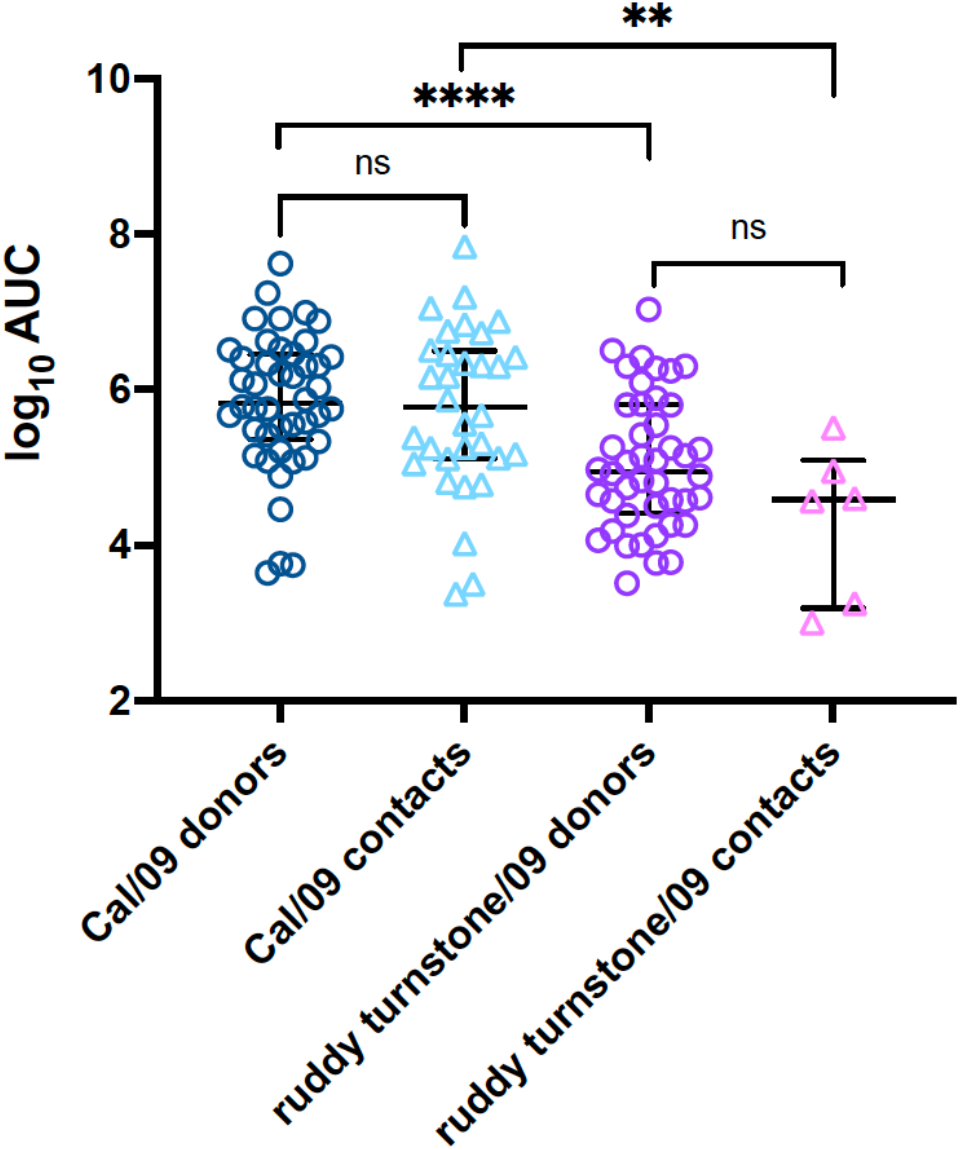
Area under the curve of infectious viral loads detected from inoculated donors or infected contacts. Data points represent AUC values from individual ferrets from which infectious virus was detected. **, p< 0.01, ****, p< 0.0001, Mann-Whitney test.

### Transmissibility of avian A(H1N1) influenza virus

We further evaluated the range of heterogeneity present in transmission results when using the A(H1N1) avian influenza virus A/ruddy turnstone/Delaware/300/2009 (ruddy turnstone/09) (18, 19), which has been reported to transmit in ferrets via respiratory droplets under the experimental setting of donor: direct contact: respiratory droplet contact at a 1:1:1 ratio, but not at a 1:1 donor:respiratory droplet contact ratio (R Fouchier, unpublished data) (18, 19). Here, the experimental setup and conditions were identical to those assessing Cal/09 virus transmissibility including a donor: respiratory droplet contact 1:1 ratio with no direct contact ferret. Transmission of an egg-derived isolate of ruddy turnstone/09 virus to exposed respiratory droplet contacts was only observed in 4 out of the 11 laboratories, with the transmission frequencies ranging from 25-75% across these four laboratories (Table 1). Compared to Cal/09 virus, there were greater differences in the ruddy turnstone/09 virus transmission outcomes across 11 laboratories, but the difference did not reach statistical significance by Fisher’s exact test of homogeneity (p=0.068). Viral shedding and seroconversion to ruddy turnstone/09 virus were detected from 6/43 exposed contact ferrets across all laboratories, resulting in a transmission efficiency of 14.0%, which was significantly lower compared to that of Cal/09 virus (72.3%, paired t-test, p < 0.001).

From the inoculated donor ferrets, the peak viral titers detected in the nasal washes or throat swabs were at 4.85 ± 0.94, mean ± SD log_10_ TCID_50_/mL after normalization, which was significantly lower than those detected in the Cal/09 inoculated donors (Mann-Whitney test, p <0.0001). Peak titers were detected from 88.6% (39/44) donors on the first sampling time point (1 or 2 dpi) followed by a decline of infectious titer over time (Figure 1B). Compared with Cal/09 virus inoculated donors, the mean ± SD log_10_ AUC of ruddy turnstone/09 virus-inoculated ferrets was 5.06 ± 1.86, significantly lower than those inoculated with the Cal/09 virus (Mann-Whitney test, p <0.0001) (Figure 2). Overall, ruddy turnstone/09 virus-inoculated donor ferrets shed lower titers of infectious virus than the Cal/09 virus-inoculated donors.

In contrast to the transmission efficiency of Cal/09 virus with a median serial interval of 3 days, for the ruddy turnstone/09 transmission experiments, the serial interval was 3 days, 5 days, or 7 days for 33.3% (2/6), 33.3% (2/6), and 33.3% (2/6) of the infected contact ferrets, respectively, with the median serial interval at 5 days. Peak viral loads (3.94 ± 0.94 mean ± SD log_10_ TCID_50_/mL) detected from the six infected contact ferrets were lower when compared to the Cal/09 infected contact ferrets (Mann-Whitney test, p=0.0022). Peak titers were detected from 16.7% (1/6), 33.3% (2/6), and 50% (3/6) infected contacts on 3 dpi, 5 dpi, and 7 dpi, respectively. Furthermore, ruddy turnstone/09 virus-infected contact ferrets shed significantly less infectious virus (4.31 ± 0.98, mean ± SD log_10_ AUC) when compared to those animals directly inoculated with Cal/09 virus (Mann-Whitney test, p=0.0033) (Figure 2). Taken together, there was a longer serial interval and lower infectious virus shed by ruddy turnstone/09 virus-exposed contact ferrets when compared to those exposed to Cal/09 virus.

### Contributing factors to ruddy turnstone/09 virus transmissibility

By standardizing the source stock virus, dose and volume of inoculation, and donor-to-contact ratio, we show that while infrequent discordant results were documented, the transmission outcomes of Cal/09 and ruddy turnstone/09 viruses independently performed by 11 laboratories were in general concordant, despite variabilities in the laboratory settings that were not standardized in the experiments (Supplemental Tables 2-4). As the transmission outcomes for the highly transmissible Cal/09 virus were more concordant than the less transmissible ruddy turnstone/09 virus, we attempted to examine if any variable, including those not standardized between laboratories, may have been associated with differences in ruddy turnstone/09 virus transmissibility results.

Univariable logistic regression was performed to first evaluate if donor viral shedding kinetics were linked to ruddy turnstone/09 virus transmission efficiency. However, examination of several parameters, including AUC (p=0.193), peak viral titer (p=0.197), and days to peak titer (p=0.473), were not statistically associated with different transmission outcomes observed between laboratories (Supplemental Table 5), indicating that differences observed between laboratories were not attributable to virological measurements.

Numerous studies have indicated a role for environmental parameters in virus transmissibility (20, 21). Room temperature was generally consistent across all groups, with means of daily recordings within 3°C for all experiments performed (20.5-23.2°C, Supplemental Table 4). In contrast, the relative humidity reported varied widely, with regard to both the range of daily readings reported during 14-day individual experiments (varying 1-60% between low and high readings) and the mean recordings over the entirety of each experiment (32.7% to 77.0%). Despite this variability, there was no statistically significant association between transmission of ruddy turnstone/09 virus and temperature, relative humidity, or absolute humidity (all p>0.3, Supplemental Table 5).

Experimental cage setups varied widely between different groups, with extensive heterogeneity present with regard to cage dimensions, airflow directionality and air changes per hour, distance between cages, and other parameters (Supplemental Table 3). Groups employing caging with airflow directionality from inoculated to contact cages more frequently reported moderate to high transmissibility (≥50%) of both viruses compared with groups lacking this airflow directionality (6/6 vs 3/5 groups for Cal/09 virus, 3/6 vs 1/5 groups for ruddy turnstone/09 virus), however these findings did not reach statistical significance (both p>0.3, Supplemental Table 5). Other specific features of cage setups, including distance between inoculated and contact cages and air changes per hour (ACH) were also not statistically linked to the ruddy turnstone/09 transmission outcomes (both p>0.4, Supplemental Table 5). Taken together, despite substantial heterogeneity in numerous non-standardized parameters in experimental setups employed between groups, no one feature was identified as modulating transmission outcomes to a significant degree.

### Contributing factors to virus pathogenicity

All ferrets inoculated with either Cal/09 or ruddy turnstone/09 were productively infected, however measurements of morbidity varied between groups for both viruses. Among Cal/09 virus-inoculated ferrets, mean maximum weight loss and peak rise in body temperature between groups ranged from <1.0-15.6% and 0.6-2.1°C, respectively (Supplemental Table 6, Supplemental Figure 3). Following ruddy turnstone/09 virus inoculation, infected ferrets generally exhibited greater mean maximum weight loss (up to 19.6%) and transient fevers (up to 3°C) (Supplemental Table 7, Supplemental Figure 4) compared to ferrets with Cal/09 virus infections; ruddy turnstone/09-inoculated ferrets reached humane experimental endpoints in 2/11 groups. The coefficient of variation between mean maximum weight loss reported between groups was generally similar (56% and 52% for Cal/09 and ruddy turnstone/09 viruses, respectively). No commonality with increased morbidity and ferret vendor, gender, or pre-inoculation body weight was identified. Furthermore, no association was found between morbidity and viral load (peak titer or AUC) or other environmental parameters, with the exception of room temperature (with higher mean room temperatures associated with greater mean weight loss) (Supplemental Table 8).

### Confidence in virus transmission results generated from multiple laboratories

Collectively, the results from this exercise demonstrate a capacity for groups possessing differences in facilities designs and experimental protocols to report varying levels of relative transmissibility and pathogenicity following inoculation of ferrets with the same virus. To illustrate how confidence in risk assessments of virus transmissibility can increase as results from multiple groups are combined, we evaluated the hypothetical risk of a virus capable of moderate to high transmission (defined as p ≥50% transmission events per total pairs of ferrets as defined in Table 2) or non-transmissible (defined as p≤25% transmission events). In these analyses, concordant results are defined as multiple groups identifying a virus exhibiting the same transmission capacity, and discordant results are defined as multiple groups identifying a virus with different transmission capacities, as defined above. By assuming concordant results across laboratories which permits pooling of all transmission outcomes, as few as three groups (12 pairs of ferrets) will yield a probability of over 80% to conclude moderate to high transmissibility when transmission was observed in at least half of all experiments, and a probability of over 85% to conclude low transmissibility when at most one transmission event was observed over all experiments.

**Table 2.**
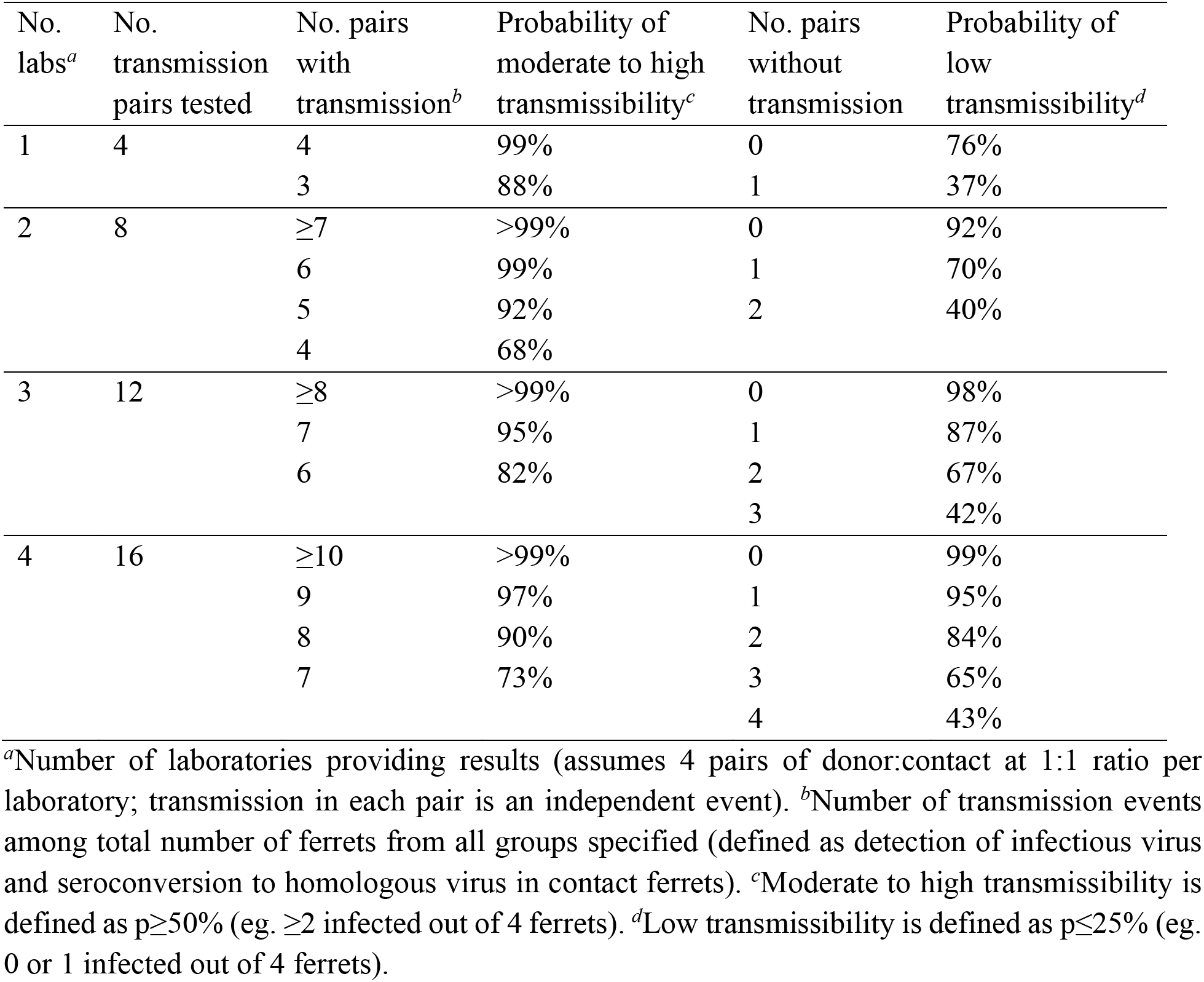
Confidence in conclusions derived from pooled samples from multiple laboratories.

Alternatively, a voting system can be considered by first drawing a conclusion on transmissibility in each laboratory, with an overall conclusion drawn based on these ‘votes’ from multiple labs. When testing for moderate to high transmissibility, and assuming n=4 ferrets per laboratory, 3 laboratories are needed to conclude moderate to high transmissibility with confidence >90% if concordant results are obtained. In agreement with probabilities shown in Table 3, a greater number of laboratories contributing results are needed to demonstrate statistically significant results when testing for low transmissibility; to conclude low transmissibility with >90% confidence, this would necessitate 5 contributing laboratories if concordant results are obtained. In this scenario, a greater number of contributing laboratories (or a greater number of donor:contact pairs per laboratory) would be required if the true transmission probability was higher for confirming low transmissibility, or when the true transmission probability was lower for confirming moderate to high transmissibility.

**Table 3.**
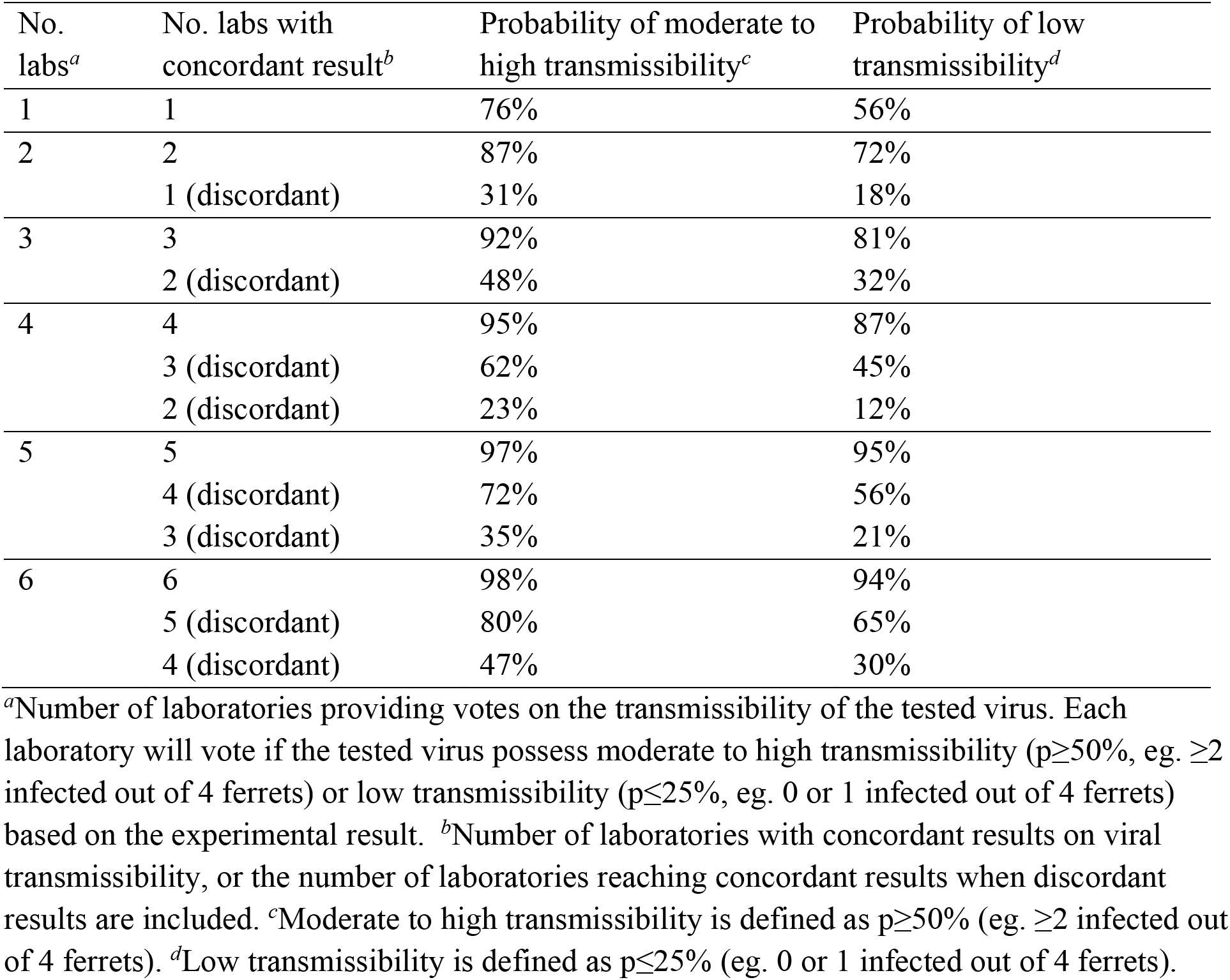
Confidence in conclusions derived from multiple laboratories considering a voting system.

Despite generally consistent results between all groups in this exercise, discordant results are possible (Table 1), highlighting the need to better understand how to responsibly interpret and account for these findings. As such, we also considered the scenario when discordant results between laboratories are recorded. To demonstrate moderate to high transmissibility, we found that 6 laboratories with 1 discordant result could still provide 80% confidence in the conclusion, while any discordant result significantly reduced confidence for concluding low transmissibility (Table 3). In both scenarios, if the results from different laboratories were more heterogeneous, the uncertainty around the conclusion from each lab increases and the overall confidence would decrease. This exercise is an illustration of the possible scenarios and confidence in drawing conclusions on transmissibility but would be affected by how moderate to high or low transmissibility were defined.

## Discussion

The importance of the ferret model for influenza virus risk assessment studies cannot be understated (4, 22). Recent advances in molecular biology, aerobiology, genomics, and other areas highlight the ways the ferret model in general, and studies evaluating virus transmissibility by the airborne route specifically, continue to contribute towards our understanding of influenza viruses and the threat they pose to human health (23–25). However, as this model becomes more commonly employed in laboratories worldwide, there is a pressing need to capture the level of variability and heterogeneity intrinsic to this research. Cross-laboratory exercises have been employed in the past to evaluate the reproducibility of assays employed for influenza virus public health efforts (26), but no such exercise has been performed to date evaluating influenza virus transmissibility in the ferret. In this exercise, 11 laboratories across different continents independently evaluated the transmission potential of Cal/09 and ruddy turnstone/09 viruses with distinct transmission potential. With only a few experimental parameters (common virus stock, standardized inoculation dose, route, volume, and the 1:1 donor:contact ratio) being controlled across the participating laboratories, we observed homogenous transmission outcomes (that is, outcomes did not differ statistically) across laboratories. Our results demonstrate the robustness of the ferret model in influenza risk assessment studies.

Risk assessment rubrics have thoroughly evaluated a wide scope of influenza A viruses, from viruses associated with poultry outbreaks in the absence of confirmed human infections, to viruses such as A(H5N1) and A(H7N9) influenza viruses that have caused substantial human disease and death (3, 27). As such, there is a need to evaluate heterogeneity of ferret transmission models employing viruses possessing a similar scope of transmissibility phenotypes. While the variability in transmission results for either the Cal/09 or ruddy turnstone/09 viruses tested in this study were not statistically significant, the range of results obtained, especially with the ruddy turnstone/09 virus, nonetheless illustrates a level of variability that can be present in transmission readouts of viruses exhibiting both low to high transmission efficiency (Table 1). This variability was present despite a high degree of standardization of virus stock, inoculation procedures, and uniformity of donor:contact ratio.

As shown in the Supplemental Methods and Supplemental Tables 1-6, this exercise captured the extensive heterogeneity in laboratory protocols and setups present between different groups. Documented variation was present in every parameter examined, inclusive of ferrets, cage setups, titration methods, and environmental conditions, among other features. Caging and airflow considerations were especially variable (Supplemental Table 2). It is impossible to standardize all contributing variables to these experiments, as institutional, animal welfare, and governmental guidelines and requirements vary worldwide, as do cost implications. That said, this exercise supports the capacity to harmonize results generated between disparate groups when a small number of procedural parameters are fixed. Interestingly, the four groups that detected infectious virus in contact nasal wash specimens in ruddy turnstone/09 transmission experiments all found 4/4 virus transmission in the Cal/09 experiment; transmission percentages between the two viruses were highly correlated between laboratories (Spearman correlation = 0.86, p < 0.001). Furthermore, while directional airflow (OR=4) did not reach statistical significance, it is nonetheless of note that 3/4 laboratories for which ruddy turnstone/09 virus transmission was detected possessed directional airflow, versus 3/7 of the laboratories for which transmission with this virus was not detected; directional airflow from inoculated to contact animals was a feature in 6/11 laboratories in this exercise (Supplemental Table 3). While our results did not conclusively identify any one experimental parameter statistically associated with enhanced transmissibility outcomes, it is possible that a confluence of parameters is nonetheless capable of creating a more permissive environment for virus transmission to occur.

To improve interpretation of results from this standardization exercise, we concurrently investigated the hypothetical confidence in concluding low transmissibility (≤25% or ≤1 ferret infected out of 4 ferrets) or moderate to high transmissibility (≥50% or ≥2 ferrets infected out of 4 ferrets) from multiple contributing laboratories. These analyses assumed both a uniform prior distribution for the transmission probability for a novel pathogen, and independent transmission outcomes from the laboratories. We considered two scenarios: one scenario where strong homogeneity across laboratories could be assumed so the samples were pooled from multiple laboratories, and another scenario where each laboratory drew their own conclusion on transmissibility such that an overall conclusion was drawn as a voting system. As influenza viruses of notable public health importance are frequently assessed across multiple independent laboratories, these analyses provide a framework to rigorously interpret independently generated findings, especially when discordant results between laboratories are reported. This is most critical in the event of a novel virus believed to possess moderate-to-high transmissibility; our analyses support that 4 independent laboratories with concordant results supporting an enhanced transmissibility phenotype yields a 95% probability of this finding, with additional independent groups or a greater number of total ferret donor:contact pairs necessary when discordant results are present.

Collectively, the findings of this exercise support the potential benefit of increased uniformity, or standardization, of some parameters when conducting risk assessment-specific activities on the same viruses. Specifically, the donor:contact ratio represents such a parameter. For a virus with moderate to high transmissibility, such as Cal/09 virus, modulation of this ratio (e.g., conducting experiments with a 2:1 donor:contact ratio, as is the case when transmission evaluations in a direct contact setting and via respiratory droplets employ a common donor) would not substantially alter conclusions drawn. However, for a virus with reduced transmissibility at a 1:1 ratio, such as the ruddy turnstone/09 virus evaluated here, it is likely that an increased donor:contact ratio (eg., 2:1) may enhance transmissibility by increasing virus-laden aerosols exhaled from infected ferrets. Previous studies on ruddy turnstone/09 virus demonstrated airborne transmission potential when employing a donor: direct contact: aerosol contact at 1:1:1 ratio; efficient transmission by direct contact will subsequently affect the quantity and kinetics of virus-laden aerosols that mediate transmission by air (18, 19). There is a need to better understand how modulation of this ratio contributes to assessments of virus transmissibility. However, this does underscore the potential complications posed by harmonizing data generated for risk assessment purposes for which the donor:contact ratio diverges. With increased heterogeneity in results between labs, uncertainty around the conclusions increases, and there is a corresponding decrease in confidence in the results (Table 3), showing the utility in increasing homogeneity across findings from different labs in order to reduce the total number of labs required to yield statistically meaningful results in this sort of analysis.

The emergence of SARS-CoV-2 further corroborates the pandemic potential of viruses of zoonotic origin. Early identification and risk assessments of novel viruses are essential for preventing the next pandemic. Continued optimization and refinement of risk assessment protocols will facilitate data interpretation in response to emerging pandemic threats. Collectively, a greater appreciation of this heterogeneity, and understanding of the scope of variability present in risk assessment settings, will permit more robust conclusions to be drawn from these efforts in the future.

## Materials and methods

### Viruses

The A(H1N1)pdm09 virus A/California/07/2009 (Cal/09) was propagated in MDCK cells (passage C3) at the US CDC as described previously (28). The low pathogenic avian influenza A(H1N1) virus A/ruddy turnstone/Delaware/300/2009 (ruddy turnstone/09) was propagated in eggs (passage E3) by St. Jude Children’s Research Hospital as described previously (19). Stocks were fully sequenced and tested for exclusivity to rule out the presence of other influenza virus subtypes prior to distribution.

### Animal and experimental variability

Groups obtained ferrets from multiple vendors and independent breeders from North America, Europe and Asia, and animals varied in their age, gender, health status, and other parameters (Supplemental Table 1). There was substantial differences between laboratories in the specific caging employed for transmission experiments, distance between cages, airflow directionality between cages, and air changes per hour (Supplemental Table 2). Anesthesia protocols, sample collection methods, and decontamination procedures to prevent cross-contamination between contact and donor animals varied between groups and are reported in Supplemental Methods. All experiments were performed under country-specific legal guidelines and approved institutional-specific animal protocols as specified in the Supplemental Methods.

### Standardized procedures

All laboratories received common stock viruses prepared by CDC and St. Jude Children’s Research Hospital with the shipping temperature recorded. Stock viruses were diluted to 10^6^ plaque forming units (PFU) in 500μl PBS based on predetermined viral titers, and donor ferrets were inoculated intranasally under in-house protocols for anesthesia (Supplemental Methods). On day 1 post-inoculation, one respiratory droplet contact ferret was introduced and exposed to each donor by housing in an adjacent cage, employing a strict 1:1 donor:contact ratio, with 4 transmission pairs tested for each virus. Ferret temperatures, weights, and nasal washes/swabs were collected every 24-48 hours. Daily room temperature and relative humidity readings were collected and are reported in Supplemental Table 3 employing pre-validated thermohygrometers with comparable readings (Testo Inc., 608-H1). Sera was collected at the end of each experiment for determination of seroconversion to homologous virus by hemagglutinin inhibition assay using established in-house serology protocols.

### Sample titration and normalization

Infectious virus titers were determined by plaque assay, 50% tissue culture infectious dose (TCID_50_) assay, or 50% egg infectious doses (EID_50_) assay at each laboratory with varying limits of detection (Supplemental Table 4). To facilitate subsequent statistical assessments across laboratories, reported titers from each laboratory were normalized to TCID_50_/mL for each virus based on PFU, TCID_50_, and EID_50_ values pre-determined by a single laboratory to minimize titration methodology-specific variation.

### Data blinding and analyses

Data blinding, aggregation and all statistical analyses were performed by an independent statistician. Transmission outcomes were compared across laboratories by each virus, using Fisher’s exact test of homogeneity. Viral load between viruses were compared by testing difference in area under the curve (AUC) using t-test. Factors associated with transmissibility and morbidity were assessed by using logistic regression and linear regression models. We also investigated the confidence in concluding low transmissibility (≤25%, or ≤1 ferret infected out of 4 ferrets) or moderate to high transmissibility (≥50% or ≥2 ferrets infected out of 4 ferrets) from multiple contributing laboratories. We assumed a uniform prior distribution for the transmission probability for a novel pathogen was assumed, and independent transmission outcomes from the laboratories. The confidence of drawing conclusion on transmissibility with concordant or discordant outcomes from the laboratories is presented. We considered a scenario where strong homogeneity across laboratory can be assumed so the samples were pooled from multiple laboratories, and another scenario that each laboratory draw their own conclusion on transmissibility and the overall conclusion was drawn as voting system. All analyses were conducted in R version 4.0.4 (R Development Core Team).

## Acknowledgements

This study was supported by Contract HHSN272201400006C from NIAID, NIH, USA. The findings and conclusions in this report are those of the authors and do not necessarily reflect the views of the Centers for Disease Control and Prevention/Agency for Toxic Substances and Disease Registry.

