## Supplemental Methods for "Robustness of the ferret model for influenza risk assessment studies: a cross-laboratory exercise"

**Supplemental Methods for Group-Specific Ferret Sample Collection Parameters.**

Ethics statement. All experiments were performed under country-specific legal guidelines and approved institutional-specific animal protocols as follows (institutes are listed by alphabetical order): Centers for Disease Control and Prevention under IACUC approved protocol 3075MAIFERC; Erasmus MC under license number AVD101002015340 and permit number 15-340-16; Harbin Veterinary Research Institute of the Chinese Academy of Agricultural Sciences under approved protocol HPAIVCZ-2-2-2017; Imperial College London, approved by the local genetic manipulation (GM) safety committee, St. Mary's Campus (center number GM77), and the Health and Safety Executive of the United Kingdom. Animal research was carried out under a United Kingdom Home Office License, P48DAD9B4; Pennsylvania State University Institutional Animal Care and Use Committee (IACUC) under protocol No. 201800250; St Jude Children's Research Hospital under IACUC approved protocol 0428; University of Georgia under IACUC approved protocol #A2019 03-021-Y3-A6; The University of Hong Kong under Committee on the Use of Live Animals in Teaching and Research approved protocol No. 3351-14; The University of Melbourne Biochemistry & Molecular Biology, Dental Science, Medicine, Microbiology & Immunology, and Surgery Animal Ethics Committee, under approval number AEC#1714278, in accordance with the NHMRC Australian code of practice for the care and use of animals for scientific purposes (8th edition); University of Pittsburgh in compliance with the guidelines of the IACUC (approved protocols #16077170 and #19075697); University of Wisconsin-Madison under IACUC approved protocol V806.

Group A: Ferrets were anesthetized for all procedures with an intramuscular (i.m.) injection (0.2-0.5 ml) of a ketamine cocktail (25 mg/kg Ketamine, 0.05 mg/kg Atropine, 2 mg/kg Xylazine) in the hamstring. To monitor subcutaneous temperatures, a subcutaneous temperature transponder, 14mm x 2mm in size (IPTT-300, BMDS, Seaford, DE) was inserted into the dorsal space between the scapulae. Nasal wash samples were obtained by introducing a 1ml volume of PBS into the nasal passages to induce sneezing, and collecting the aspirate using a sterile plastic dish. 70% EtOH surface decontamination of gloves and equipment was performed between handling contact animals; contact animals always handled prior to virus-inoculated animals.

Group B: Ferrets were anesthetized for all procedures with an i.m. injection (0.2-0.3 ml) of a ketamine cocktail (20 mg/kg Ketamine, 1 mg/kg Xylazine) in the hamstring. A digital thermometer was employed to measure rectal temperatures. Nasal wash samples were obtained by administering a 2ml volume of PBS through the choanae via an animal feeding needle with a bent tip, connected to a 2ml syringe, collecting the PBS passively flowing out through the nostrils using a sterile plastic dish. 75% EtOH surface decontamination of gloves and equipment was performed between handling contact animals; contact animals always handled prior to virus-inoculated animals. Different utensils/tools used between virus groups; each contact ferret had a dedicated tool set.

Group C: Ferrets were anesthetized for all procedures with an i.m. injection (0.5 ml/kg) of a ketamine cocktail (20 mg/kg Ketamine, 1 mg/kg Xylazine) in the thighs. Temperatures were monitored daily using a subcutaneous implantable temperature transponder (Bio Medic Data Systems), located on the back of the neck. Nasal wash samples were obtained by introducing a 1ml volume of PBS into the nasal passages to induce sneezing, collecting the aspirate using a sterile plastic dish, and washing the dish with an additional 1ml of PBS. 70% EtOH surface decontamination of gloves and equipment was performed between handling contact animals; contact animals always handled prior to virus-inoculated animals. Different bite-resistant gloves, utensils and tools used between contact and virus-inoculated animals, with disposable gowns and gloves changed between isolators.

<u>Group D</u>: Ferrets were anesthetized for all procedures with an i.m. injection (0.04-0.1 ml) of a mixture of Ketamine (4-5 mg/kg) and Dexmedetomidine (10-40 μg/kg) in the shoulder, antagonized with a 1:1 volume of Atipamezole (0.05-0.2 mg/kg). To monitor subcutaneous temperatures, a transponder chip (IPTT-300, BMDS) was implanted in the back of the neck and scanned by a portable reader. Nasal wash samples were obtained by administering a 2ml volume of PBS through the choanae via an animal feeding needle with a bent tip, connected to a 3ml syringe, collecting the PBS passively flowing out through the nostrils. 70% EtOH surface decontamination and changing of gloves and sleeves was performed between handling contact animals; contact animals always handled prior to virus-inoculated animals. Different containers and utensils/tools used between virus groups.

<u>Group E</u>: Ferrets were anesthetized for virus inoculation with an i.m. injection (0.2-0.3 ml) of a ketamine cocktail (100 mg/ml Ketamine, 20 mg/ml Xylazine), and for nasal wash sample collection with an i.m. injection (0.2-0.3 ml) of Xylazine (20 mg/ml), in the hind-leg. To monitor subcutaneous temperatures, a 13mm x 2.12mm transponder (Lifechip, Bio-Thermo) was implanted into the dorsal region of each ferret. Nasal wash samples were obtained by introducing a 1ml volume of PBS into the nasal passages to induce sneezing, collecting the aspirate into a 10ml

sample jar. 80% EtOH surface decontamination of gloves and equipment was performed between handling contact animals; contact animals always handled prior to virus-inoculated animals.

Group F: Ferrets were anesthetized for virus inoculation with an i.m. injection (0.2 – 0.3 ml) of a mixture of Ketamine (25 mg/kg) and Medetomidine (0.03 mg/kg), antagonized with Atipamezole (0.15 mg/kg), and for throat swabbing with an i.m. injection (0.2 – 0.3 ml) of Ketamine (20 mg/kg) in the hamstring. A digital thermometer was employed to measure rectal temperatures. Throat swabs were obtained and collected in 1ml of virus transport medium. 70% EtOH surface decontamination of gloves and equipment was performed between handling contact animals; contact animals and virus-inoculated animals were handled on alternate days.

Group G: Ferrets were anesthetized for virus inoculation with inhaled isoflurane, and for nasal wash sample collection with an i.m. injection (0.25-0.5 ml) of Ketamine (25 mg/kg), in the hamstring. Microchip transponders (Bio Medic Data Systems) were used for subcutaneous temperature monitoring. Nasal wash samples were obtained by introducing a 1ml volume of PBS into the nasal passages to induce sneezing, collecting the aspirate using a sterile plastic specimen cup. 70% EtOH surface decontamination of gloves and equipment was performed between handling contact animals; contact animals always handled prior to virus-inoculated animals. Outer PPE was changed between cubicles.

Group H: Ferrets were anesthetized for virus inoculation and euthanasia with an i.m. injection (0.25-0.5ml) of a mixture of Ketamine (22 mg/kg infection, 25 mg/kg euthanasia) and Xylazine (0.9 mg/kg); no anesthesia was employed for nasal wash collection. To monitor subcutaneous temperatures, a subcutaneous temperature transponder, 14mm x 2mm in size (IPTT-300, BMDS, Seaford, DE) was inserted into the flank. Nasal wash samples were obtained by introducing a 2ml volume of PBS into the nasal passages to induce sneezing, collecting the aspirate using a plastic cone. 70% EtOH surface decontamination of gloves and equipment was performed between handling contact animals; contact animals always handled prior to virus-inoculated animals.

Group I: Ferrets were anesthetized for virus inoculation (i.m. injection of 25 mg/kg Ketamine and 2 mg/kg Xyzaline in the hamstring) or nasal wash collection (i.m. injection of 25 mg/kg Ketamine in the hamstring). Rectal temperature was taken using separate thermometers for each animal. Nasal wash samples were obtained by introducing a 1ml volume of PBS into the nasal passages to induce sneezing, collecting the aspirate using a sterile sample cup. 1% Virkon surface decontamination was performed between handling contact animals; contact animals always handled prior to virus-inoculated animals. Different utensils/tools used between virus groups; each contact ferret had a dedicated tool set.

Group J: Ferrets were anesthetized for all procedures with an i.m. injection (0.2-0.6 ml) of a ketamine cocktail (20 mg/kg Ketamine, 0.5 mg/kg Atropine, 2 mg/kg Xylazine), in the hind leg. To monitor subcutaneous temperatures, a transponder (IPT-300, BMDS, Seaford, DE) was implanted using a 12 gauge syringe delivery system in the dorsal space between the scapulae (i.e. in the scruff). Nasal wash samples were obtained by introducing a 1ml volume of PBS into the nasal passages to induce sneezing, collecting the aspirate using a sterile plastic dish, and washing the dish with an additional 1ml of PBS. 70% EtOH or Pre-Empt surface decontamination was performed between handling contact animals; contact animals always handled prior to virus-inoculated animals. Change of lab coats and gloves was performed between each transmission pair.

Group K: Ferrets were anesthetized for all survival procedures with inhaled isoflurane. To monitor subcutaneous temperatures, a subcutaneous temperature transponder chip, 14mm x 2mm in size (IPTT-300, BMDS, Seaford, DE) is placed under the skin on the dorsal midline, usually between the scapulae. Nasal wash samples were obtained by administering ~2mL of 1x PBS to one nostril using a 3mL syringe and collecting the PBS passively flowing out through the nostrils. A concentrated cleaning solution blend (1:16 ratio of Hydrogen Peroxide 0.1-0.3% pH 2.3) was employed for surface decontamination of gloves and equipment between handling contact animals; contact animals always handled prior to virus-inoculated animals. Change of gloves and disinfection of all tools and biosafety cabinet was performed between each transmission pair.
