## Supplemental Tables and Figures for "Robustness of the ferret model for influenza risk assessment studies: a cross-laboratory exercise"

**Supplemental Table 1. Summary of titration methodology by each laboratory.**

| Group | Titers determined in each laboratory | | | | After normalization[d] ($TCID_{50}$/ml) | |
|---|---|---|---|---|---|---|
| | Units[a] | LOD/ml[b] | Cal/09[c] | Ruddy turnstone/09[c] | Cal/09 | Ruddy turnstone/09 |
| A | PFU | 10 | $9.0x10^6$ | $6.6x10^5$ | $1.8x10^7$ | $1.32x10^6$ |
| B | $EID_{50}$ | 5.62 | $2.51x10^6$ | $2.29x10^7$ | $4.46x10^6$ | $4.07x10^7$ |
| C | $TCID_{50}$ | 10 | $1.0x10^5$ | $1.58x10^6$ | $1.0x10^5$ | $1.58x10^6$ |
| D | PFU | 10 | $7.0x10^7$ | $1.65x10^7$ | $1.4x10^8$ | $3.30x10^7$ |
| E | $TCID_{50}$ | 100 | $1.86x10^6$ | $9.33x10^5$ | $1.86x10^6$ | $9.33x10^5$ |
| F | $TCID_{50}$ | 3.16 | $6.31x10^6$ | $1.0x10^7$ | $6.31x10^6$ | $1.0x10^7$ |
| G | $TCID_{50}$ | 10 | $5.0x10^6$ | $2.5x10^6$ | $5.0x10^6$ | $2.5x10^6$ |
| H | PFU | 10 | $7.0x10^6$ | $2.6x10^6$ | $1.4x10^7$ | $5.2x10^6$ |
| I | $TCID_{50}$ | 61.5 | $4.9x10^7$ | $6.0x10^7$ | $4.9x10^7$ | $6.0x10^7$ |
| J | $TCID_{50}$ | 3.16 | $1.58x10^7$ | $5.62x10^5$ | $1.58x10^7$ | $5.62x10^5$ |
| K | $TCID_{50}$ | 3.16 | $2.82x10^6$ | $1.58x10^6$ | $2.82x10^6$ | $1.58x10^6$ |

[a]PFU, plaque forming units. $EID_{50}$, 50% egg infectious dose. $TCID_{50}$, 50% tissue culture infectious dose (employing MDCK cells). [b]LOD/ml, limit of detection for titration of ferret specimens for the units specified. [c]Titration of received virus employing laboratory assay-specific methods. [d]For standardization purposes, all infectious titers were normalized to $TCID_{50}$/mL based on pre-determined titers for the Cal/09 virus ($5x10^6$ PFU/ml, $5.01x10^6$ $EID_{50}$/ml, and $1.0x10^7$ $TCID_{50}$/ml), and ruddy turnstone/09 virus ($2.5x10^6$ PFU/ml, $3.16x10^8$ $EID_{50}$/ml, and $3.16x10^6$ $TCID_{50}$/ml).

**Supplemental Table 2. Ferret source and health status prior to study.**

| Group | Commercial vendor | Spayed/neutered /descented | Gender | Vaccination status | Additional treatments |
|---|---|---|---|---|---|
| A | Triple F Farms, USA | Yes | M | rabies, distemper | none |
| B | Independent breeders | Yes | F | distemper, parvovirus | none |
| C | Triple F Farms, USA | Yes | F | rabies, distemper | meloxicam, penicillin, ivermectin, ponazril |
| D | Triple F Farms, USA | Yes | F | rabies, distemper | meloxicam, penicillin, ivermectin, ponazril, |
| E | Independent breeders | Yes (M only) | M/F | distemper | hormone blocker (F only) |
| F | Triple F Farms, USA | Yes | F | rabies | none |
| G | Triple F Farms, USA | Yes | M | rabies, distemper | none |
| H | Highgate Farms, UK | No | F | none | delvosteron (F only) |
| I | Independent breeders | Yes | M | rabies, distemper, parvovirus | none |
| J | Triple F Farms, USA | Yes | M/F | rabies, distemper | none |
| K | Triple F Farms, USA | Yes | M | rabies, distemper | none |

**Supplemental Table 3. Caging and physical environment.**

| Group | cage dimensions LxWxH (cm)[a] | cage material | cage interface distance | transmission interface | airflow to contacts[b] | ACH[c] |
|---|---|---|---|---|---|---|
| A | 56 x 42 x 42 | stainless steel | 0.3-1 cm | two perforated side-walls with openings <5 mm | no | 150-180 |
| B | 25 x 33 x 40 | stainless steel | 4 cm | double-layered net divider | yes | 20-50 |
| C | 36 x 36 x 41 | stainless steel | 5 cm | two perforated side-walls with openings 13 mm | no | 12-15 |
| D | 28 x 38 x 30 | stainless steel | 5 cm | stainless steel mesh | no | 45 |
| E | 28.3 x 78.7 x 41 | stainless steel | 2 cm | double-layered stainless steel perforated divider with openings 5 mm | yes | 36-44 |
| F | 50 x 30 x 30 | perspex | 10 cm | perforated stainless steel grid with openings 10 mm | yes | 30-40 |
| G | 72 x 61 x 41 | stainless steel | 0.5-1 cm | two perforated side-walls with openings 5 mm | no | 15 |
| H | 78 x 50 x 50 | perspex/stainless steel | 0.25 cm | perforated stainless steel grid with openings 5 mm | no | 15 |
| I | 61 x 61 x 48 | stainless steel | 4.5-5 cm | perforated stainless steel grid with openings 5 mm | yes | 39-41 |
| J | 68.6 x 28 x 40.6 | stainless steel | 2.5 cm | two perforated side-walls with openings 5 mm | yes | 25 |
| K | 28 x 79 x 41 | stainless steel | 2.5 cm | two perforated panels with staggered openings 5mm | yes | 25-35 |

[a]Housing environment of individually-housed ferrets. [b]Yes, directional airflow from inoculated to contact cages; No, airflow is ambient or otherwise not directional from inoculated to contact cages. [c]ACH, air changes per hour within primary containment area.

**Supplemental Table 4. Environmental summaries for transmission experiments.**

| Group | Virus | temperature (°C) | | relative humidity (% RH) | |
|---|---|---|---|---|---|
| | | mean | range | mean | range |
| A | Cal/09 | 21.1 | 21.0-21.2 | 54.1 | 53.5-55.0 |
| | Ruddy turnstone/09 | 21.0 | 20.2-21.2 | 54.6 | 51.9-56.7 |
| B | Cal/09 | 21.4 | 21.2-22.8 | 32.9 | 30.2-38.5 |
| | Ruddy turnstone/09 | 21.4 | 21.2-22.8 | 32.9 | 30.2-38.5 |
| C | Cal/09 | 22.2 | 20.7-24.1 | 77.0 | 61.0-86.5 |
| | Ruddy turnstone/09 | 22.0 | 21.0-23.4 | 67.5 | 39.9-99.6 |
| D | Cal/09 | 23.2 | 23.0-24.1 | 45.0 | 42.7-46.6 |
| | Ruddy turnstone/09 | 23.2 | 23.0-24.1 | 45.0 | 42.7-46.6 |
| E | Cal/09 | 21.4 | 21.0-21.9 | 52.9 | 39.0-59.0 |
| | Ruddy turnstone/09 | 21.6 | 20.9-22.0 | 51.9 | 43.8-64.5 |
| F | Cal/09 | 22.0 | 21.1-22.3 | 44.7 | 42.0-49.3 |
| | Ruddy turnstone/09 | 21.9 | 21.4-22.2 | 45.2 | 41.2-54.7 |
| G | Cal/09 | 22.3 | 21.7-23.5 | 47.5 | 41.2-59.4 |
| | Ruddy turnstone/09 | 22.3 | 21.7-23.5 | 47.5 | 41.2-59.4 |
| H | Cal/09 | 22.2 | 20.7-24.7 | 63.8 | 58.9-73.1 |
| | Ruddy turnstone/09 | 21.1 | 20.6-21.8 | 69.8 | 64.8-76.7 |
| I | Cal/09 | 20.5 | 20.2-21.1 | 55.3 | 37.7-67.0 |
| | Ruddy turnstone/09 | 20.5 | 20.2-21.1 | 55.3 | 37.7-67.0 |
| J | Cal/09 | 22.9 | 22.1-23.6 | 40.8 | 36.0-49.4 |
| | Ruddy turnstone/09 | 21.7 | 20.8-23.2 | 32.7 | 32.0-33.0 |
| K | Cal/09 | 21.9 | 21.2-22.3 | 47.8 | 45.1-54.9 |
| | Ruddy turnstone/09 | 21.8 | 21.3-22.2 | 48.3 | 43.8-59.1 |

Pre-validated hygrometers with comparable readings were employed to measure temperature and relative humidity in each facility; hygrometers were pre-calibrated at a central location prior to distribution to all participating groups. All groups employ a 12-hour light/dark cycle with the exception of Group E, which employs a 9-hour light/dark cycle.

**Supplemental Table 5. Parameters associated with transmission[a] of ruddy turnstone/09 by univariable logistic regression.**

| **Parameters[b]** | **Unadjusted OR (95% CI)** | **p-value** |
|---|---|---|
| Donor viral load ($log_{10}$ AUC)[c] | 0.18 (0.01, 2.37) | 0.193 |
| Donor peak titer ($log_{10}TCID_{50}$/ml) | 0.56 (0.23, 1.35) | 0.197 |
| Donor time to peak titer (dpi) | 0.28 (0.01, 8.88) | 0.473 |
| Air change (per 10 ACH) | 0.61 (0.19, 1.96) | 0.408 |
| Directional airflow to contacts (reference: without directional airflow) | 4.00 (0.27, 60.32) | 0.317 |
| Temperature (per 0.1°C) | 1.01 (0.85, 1.21) | 0.874 |
| Relative humidity (%) | 0.94 (0.83, 1.06) | 0.318 |
| Absolute humidity (per g/$m^3$) | 0.71 (0.36, 1.42) | 0.337 |
| Distance between cages (cm) | 1.01 (0.63, 1.60) | 0.983 |

[a]Outcome measure: higher transmissibility was defined as ≥25% for A/ruddy turnstone/Delaware/300/2009. [b]All parameters, except "directional air flow to contacts" were numeric. [c]Statistically insignificant for viral load (AUC) with P-value = 0.317

**Supplemental Table 6. Clinical signs of donor ferrets following inoculation with Cal/09 virus.**

| Group | Gender | Age (months) | Body weight changes[a] | | | Temperature[a] | | | Respiratory signs[e] | Lethality[f] | RII[g] |
|---|---|---|---|---|---|---|---|---|---|---|---|
| | | | Baseline (g)[b] | Mean max loss (%)[c] | Range (day)[d] | Baseline (°C)[b] | Mean max rise (°C)[c] | Range (day)[d] | | | |
| A | M | 8 | 1465 | 8.5 | 6-9 | 38.2 | 1.8 | 1-6 | 2/4 | 0/4 | 1.00 |
| B | F | 4-6 | 961 | 5.0 | 6 | 38.3 | 0.7 | 2 | 4/4 | 0/4 | 1.27 |
| C | F | 5 | 688 | 16.9 | 3-12 | 39.7 | 0.6 (3/4) | 2-11 | 4/4 | 0/4 | 1.66 |
| D | F | 5 | 699 | 15.6 | 6-11 | 37.2 | 2.1 | 5-14 | 0/4 | 0/4 | 1.00 |
| E | M/F | 4 | 862 | 4.9 | 4-11 | 37.8 | 1.9 | 1-4 | 4/4 | 0/4 | 1.04 |
| F | F | 6-12 | 829 | 9.3 | 3-7 | 38.4 | 1.0 | 1-5 | 2/4 | 0/4 | 1.39 |
| G | M | 3-5 | 1013 | 8.6 | 1-6 | 38.4 | 1.0 | 1-4 | 3/4 | 0/4 | 1.21 |
| H[h] | F | 4-5 | 905 | 3.9 (3/4) | 2-6 | 38.2 | 0.7 | 2-10 | 3/4 | 0/4 | 1.05 |
| I | M | 4-6 | 1258 | 1.0 (1/4) | 2 | 39.1 | 0.7 | 2 | 4/4 | 0/4 | 1.07 |
| J | M/F | 6 | 1144 | 11.2 | 5-6 | 39.0 | 0.8 | 1-10 | 0/4 | 0/4 | 1.30 |
| K | M | 6-8 | 1399 | 11.5 | 6-11 | 38.1 | 1.6 | 2-5 | 1/4 | 0/4 | 1.04 |

[a]Ferret data are inclusive of n=4 unless otherwise specified. Data are reflective of measurements collected every 24 hrs (Groups A, C, D, E, G, H, K) or 48 hrs (Groups B, F, I, J). [b]Mean pre-inoculation body weight (in grams) or temperature (in °C). [c]Percentage mean maximum weight loss or mean maximum rise in temperature (in °C) (compared to baseline on day 0), detected between days 1-14 post-inoculation. Data is inclusive of all ferrets for which weight loss/temperature increases were detected during the observation period; the number of ferrets included in this mean is specified when this is not n=4. [d]Day range of maximum weight loss values reported among ferrets included in the reported mean. [e]Number of ferrets for which respiratory signs (sneezing, coughing, heavy breathing, open mouth breathing, or nasal discharge) were observed between days 1-14 post-inoculation at least once. [f]Number of ferrets that reached humane euthanasia endpoints (day of death specified in parentheses). [g]RII, relative inactivity index. [h]Reported values for this group span days 0-10 post-inoculation.

**Supplemental Table 7. Clinical signs of donor ferrets following inoculation with ruddy turnstone/09 virus.**

| Group | Gender | Age (months) | Body weight[a] | | | Temperature[a] | | | Respiratory signs[e] | Lethality[f] | RII[g] |
|---|---|---|---|---|---|---|---|---|---|---|---|
| | | | Baseline (g)[b] | Mean max loss (%)[c] | Range (day)[d] | Baseline (°C)[b] | Mean max rise (°C)[c] | Range (day)[d] | | | |
| A | M | 9 | 1415 | 10.7 | 2-11 | 38.5 | 2.0 | 1-4 | 1/4 | 0/4 | 1.09 |
| B | F | 4-6 | 1005 | 1.2 | 4-8 | 38.4 | 0.5 | 2 | 3/4 | 0/4 | 1.13 |
| C | F | 5 | 628 | 12.9 | 1-8 | 39.2 | 0.6 (3/4) | 1-2 | 3/4 | 0/4 | 1.59 |
| D | F | 5 | 678 | 17.8 | 6-12 | 37.3 | 3.0 | 2 | 0/4 | 2/4 (12) | 1.31 |
| E | F | 4 | 809 | 11.4 | 2-9 | 38.3 | 1.5 | 1-2 | 4/4 | 0/4 | 1.13 |
| F | F | 6-12 | 708 | 15.0 | 3-11 | 38.9 | 1.4 (3/4) | 1-5 | 2/4 | 1/4 (7) | 1.39 |
| G | M | 3-5 | 1050 | 16.7 | 2-5 | 37.7 | 1.9 | 2-5 | 2/4 | 0/4 | 1.27 |
| H[h] | F | 4-5 | 838 | 7.4 | 5-8 | 38.4 | 0.8 (3/4) | 2-10 | 4/4 | 0/4 | 1.73 |
| I | M | 4-6 | 1220 | 2.2 (2/4) | 2 | 39.2 | 1.2 | 2-4 | 4/4 | 0/4 | 1.23 |
| J | M/F | 6 | 1111 | 12.8 | 2-10 | 38.5 | 2.3 | 1-4 | 0/4 | 0/4 | 1.09 |
| K | M | 6-8 | 1274 | 19.6 | 7-9 | 39.0 | 1.6 | 1-5 | 1/4 | 0/4 | 1.23 |

[a]Ferret data are inclusive of n=4 unless otherwise specified. Data are reflective of measurements collected every 24 hrs (Groups A, C, D, E, G, H, K) or 48 hrs (Groups B, F, I, J). [b]Mean pre-inoculation body weight (in grams) or temperature (in °C). [c]Percentage mean maximum weight loss or mean maximum rise in temperature (in °C) (compared to baseline on day 0), detected between days 1-14 post-inoculation. Data is inclusive of all ferrets for which weight loss/temperature increases were detected during the observation period; the number of ferrets included in this mean is specified when this is not n=4. [d]Day range of maximum values reported among ferrets included in the reported mean. [e]Number of ferrets for which respiratory signs (sneezing, coughing, heavy breathing, open mouth breathing, or nasal discharge) were observed between days 1-14 post-inoculation at least once. [f]Number of ferrets that reached humane euthanasia endpoints (day of death specified in parentheses). [g]RII, relative inactivity index. [h]Reported values for this group span days 0-10 post-inoculation.

**Supplemental Table 8. Parameters associated with % weight changes of Cal/09 and ruddy turnstone/09 inoculated donors by univariable linear regression.**

| Parameters[a] | Cal/09 | | Ruddy turnstone/09 | |
|---|---|---|---|---|
| | **Difference (95% CI)** | **p-value** | **Difference (95% CI)** | **p-value** |
| Donor viral load ($log_{10}$ AUC)[b] | 1.6 (-2.5, 5.7) | 0.468 | -0.4 (-5.1, 4.4) | 0.878 |
| Donor peak titer ($log_{10}TCID_{50}$/ml) | 0.3 (-1.4, 2.0) | 0.739 | -0.2 (-2.1, 1.7) | 0.826 |
| Donor time to peak titer (dpi) | -2.2 (-11.7, 7.3) | 0.661 | -0.9 (-10.5, 8.6) | 0.852 |
| Gender (reference: male)[c] | -2.7 (-9.8, 4.3) | 0.472 | 1.4 (-7.9, 10.8) | 0.771 |
| Air change (per 10 ACH) | 0.1 (-0.7, 0.8) | 0.848 | 0.1 (-0.8, 1.0) | 0.816 |
| Directional airflow to contact (reference: without directional airflow) | 3.6 (-2.1, 9.2) | 0.251 | 2.7 (-4.5, 10.0) | 0.480 |
| Temperature (°C) | -4.4 (-7.3, -1.5) | 0.015 | -6.2 (-9.9, -2.5) | 0.009 |
| Relative humidity (%) | -0.1 (-0.3, 0.2) | 0.618 | 0.0 (-0.3, 0.4) | 0.897 |
| Absolute humidity ($g/m^3$) | -0.6 (-2.0, 0.7) | 0.365 | -0.2 (-2.0, 1.5) | 0.794 |
| Distance between cages (cm) | -0.4 (-1.5, 0.7) | 0.514 | -0.1 (-1.5, 1.3) | 0.884 |

[a]All parameters, except "directional airflow to contact" and gender were numeric. [b]Also insignificant for viral load (AUC) with P-values = 0.283 and 0.734 for Cal/09 and ruddy turnstone/09 respectively. [c]Excluded two laboratories which used both genders.

**Supplemental Figure 1. Transmission kinetics of A/California/7/2009 virus.** Nasal washes (all groups except Group F) or throat swabs (Group F) were sampled from donor (left bars) and aerosol contact ferrets (right bars) to determine infectious viral loads; titers are reported as $\log_{10}$ PFU/ml (Groups A, D, H), $\log_{10}$ $TCID_{50}$/ml (Groups C, E, F, G, I, J, K), or $\log_{10}$ $EID_{50}$/ml (Group B). Limit of detection for each graph is reported in Supplemental Table 1.

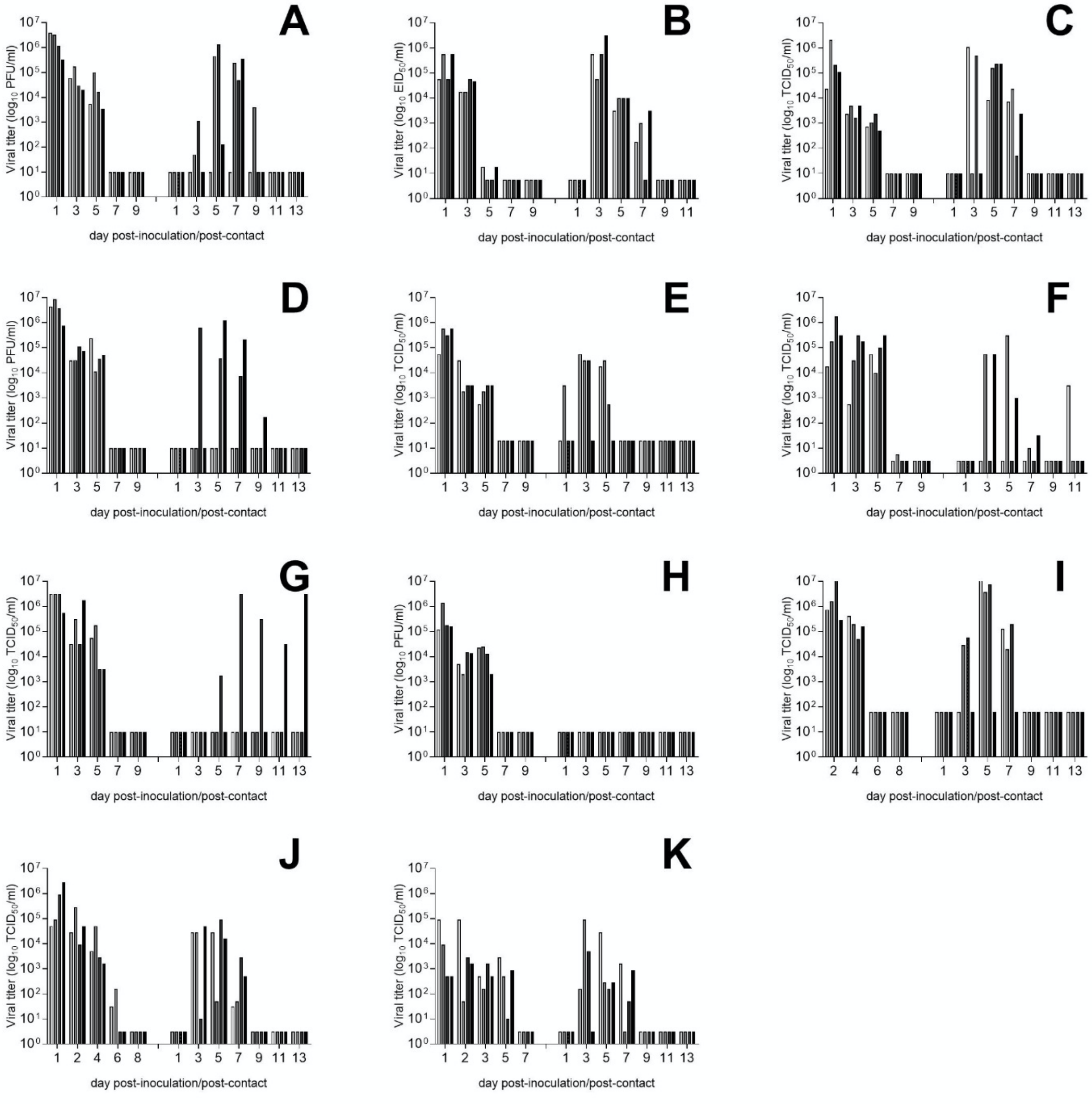

**Supplemental Figure 2. Transmission kinetics of A/ruddy turnstone/Delaware/300/2009.** Nasal washes (all groups except Group F) or throat swabs (Group F) were sampled from donor (left bars) and aerosol contact ferrets (right bars) to determine infectious viral loads; titers are reported as $\log_{10}$ PFU/ml (Groups A, D, H), $\log_{10}$ $TCID_{50}$/ml (Groups C, E, F, G, I, J, K), or $\log_{10}$ $EID_{50}$/ml (Group B). Limit of detection for each graph is reported in Supplemental Table 1.

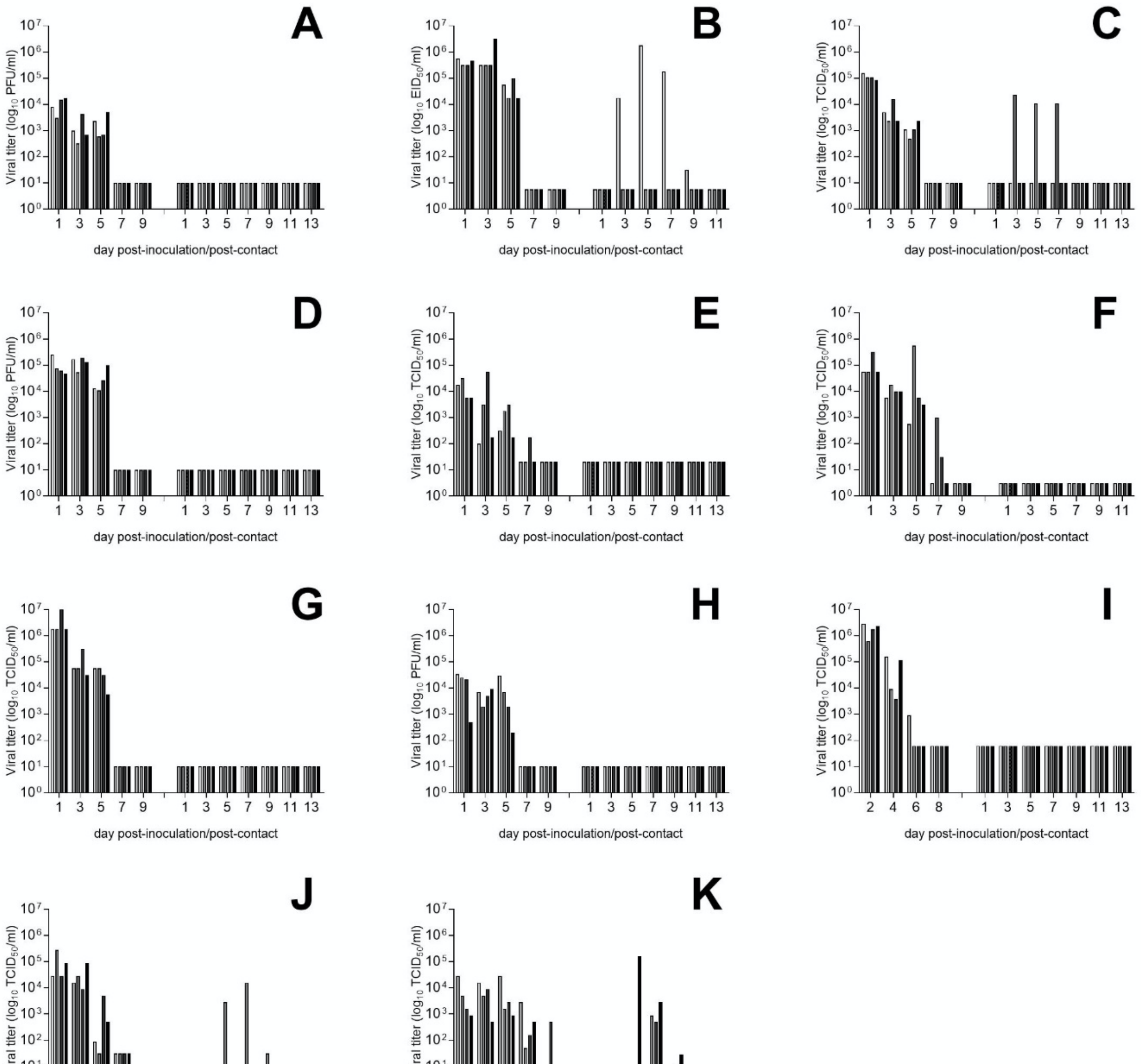

**Supplemental Figure 3. Weight loss kinetics of A/California/7/2009.** Body weights from inoculated animals were collected every day (Groups A, C, D, E, G, H, K) or every-other-day (Groups B, F, I, J) post-inoculation through the days indicated. Body weight percentages were set at 100% on the day of inoculation for each animal; lines represent individual ferrets.

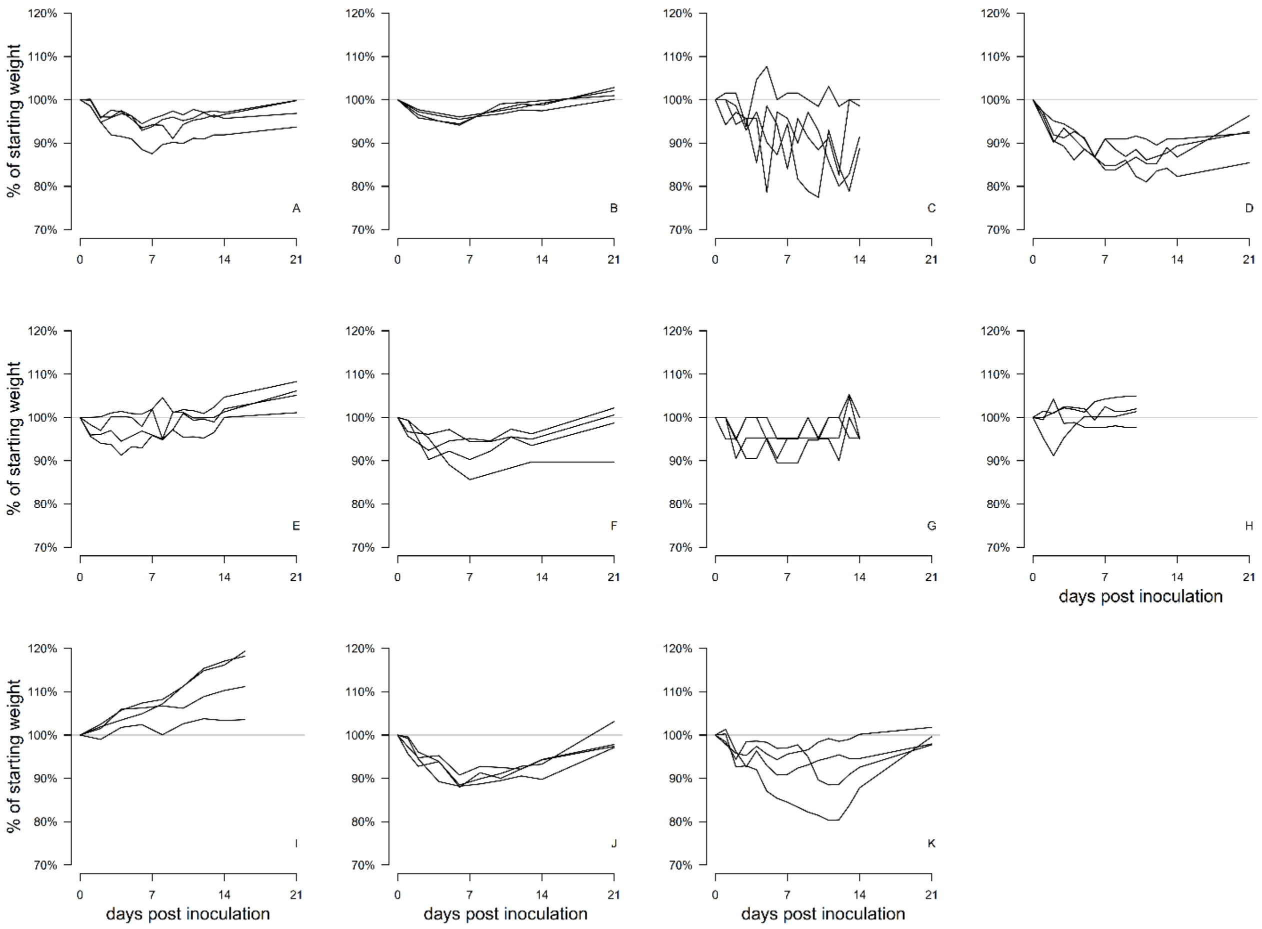

**Supplemental Figure 4. Weight loss kinetics of A/ruddy turnstone/Delaware/300/2009.** Body weights from inoculated animals were collected every day (Groups A, C, D, E, G, H, K) or every-other-day (Groups B, F, I, J) post-inoculation through the days indicated. Body weight percentages were set at 100% on the day of inoculation for each animal; lines represent individual ferrets. Group D, two ferrets were humanely euthanized day 12 p.i. due to reaching endpoint criteria. Group F, one ferret was humanely euthanized day 7 p.i. due to reaching endpoint criteria.

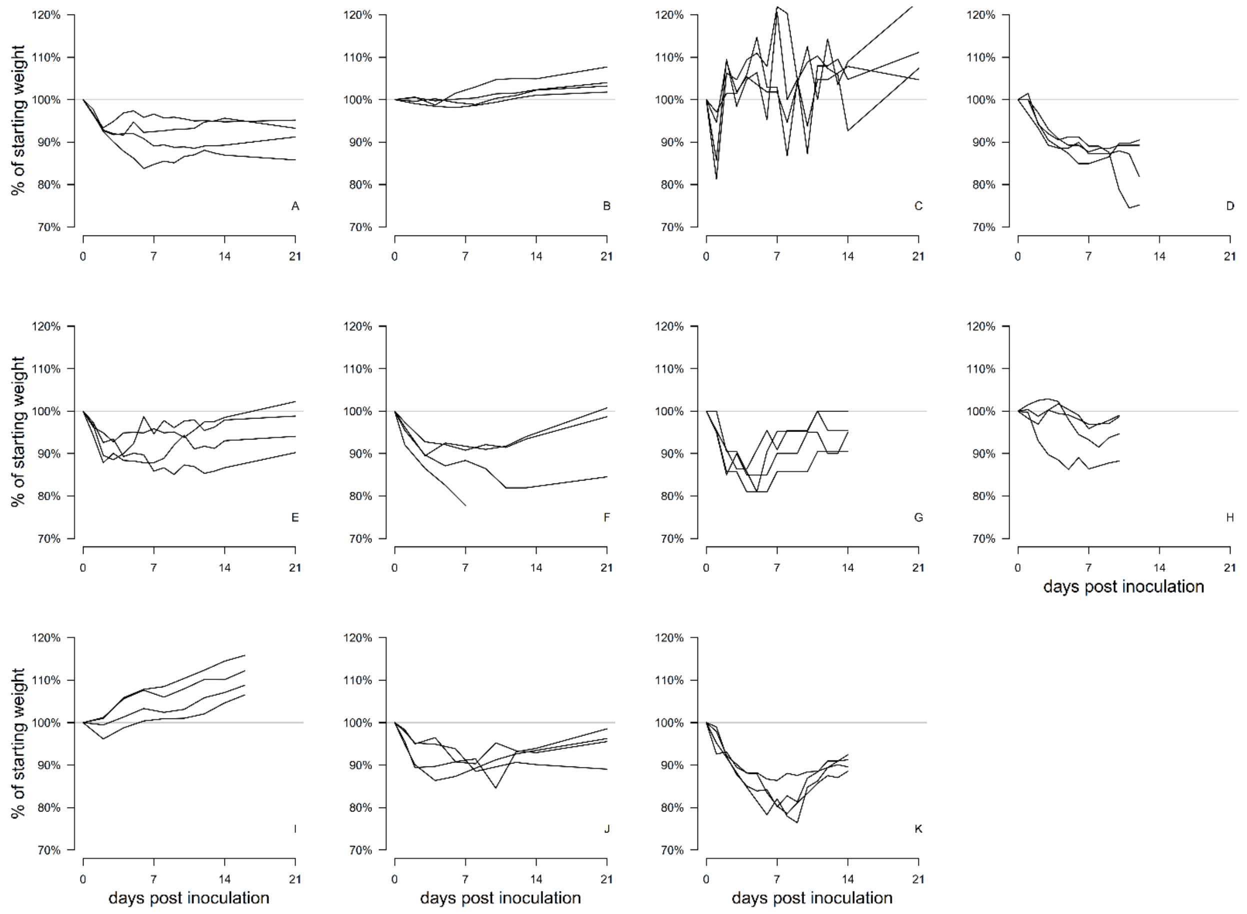
